# A whole blood intracellular cytokine assay optimised for field site studies demonstrates polyfunctionality of CD4+ T cells in acute scrub typhus

**DOI:** 10.1101/2022.10.28.513456

**Authors:** Manutsanun Inthawong, Nattapon Pinthong, Areerat Thaiprakhong, Tri Wangrangsimakul, Piyanate Sunyakumthorn, Jennifer Hill, Piengchan Sonthayanon, Daniel H. Paris, Susanna Dunachie, Barbara Kronsteiner

## Abstract

**Background:** Assessment of cellular immune responses by combining intracellular cytokine staining and immunophenotyping using flow cytometry enables the simultaneous measurement of T cell phenotype and effector function in response to pathogens and vaccines. The use of whole blood samples rather than peripheral blood mononuclear cells avoids both the need for immediate processing and loss of functional antigen presenting cells due to processing and cryopreservation. Using whole blood provides the possibility to stimulate peripheral T cells *in situ*, and is more suitable for studies where sample volume is limited, such as those involving children, the elderly and critically ill patients. The aim of this study was to provide a robust tool for the assessment of antigen-specific T cell responses in a field site setting with limited resources.

**Methodology/Principle Findings:** We optimised a flow cytometry-based whole blood intracellular cytokine assay (WBA) with respect to duration of antigen stimulation and intracellular protein retention time. We demonstrate the ability of the WBA to capture polyfunctional T cell responses in the context of acute scrub typhus infection, by measuring IFN-γ, TNF and IL-2 in CD4+ and CD8+ T cells in response to the causative agent *O. tsutsugamushi* (OT). Using an optimised OT antigen preparation, we demonstrate the presence of polyfunctional antigen-specific memory CD4+ T cells in the blood of scrub typhus patients.

**Conclusions/Significance:** In conclusion, this flow cytometry-based WBA is well-suited for use at field study sites, and enables the assessment of polyfunctional T cell responses to infectious agents and vaccines through delineation of antigen-specific cytokine secretion at the single cell level.

**Author summary:** Scrub typhus is an acute febrile illness caused by the bite of infected chigger mites transmitting *O. tsutsugamushi, a* gramnegative obligate intracellular bacteria in the family *Rickettsiaceae*. The disease progression and clinical manifestations are closely associated with host immune responses. T cell responses are in strong relation with immune protection against scrub typhus. As there is limited knowledge on specific T cell roles against scrub typhus, we optimized a flow-cytometer based protocol to assess *O. tsutsugumushi*-specific T cell responses in whole blood samples, which is suitable for studies in limited resource settings. This method requires low blood volumes, but enables multiparametric immunophenotyping assessments. The optimized WBA protocol could be a useful tool for comprehensive immunopathological studies in scrub typhus, and other infectious diseases as well as vaccine studies in areas with limited availability of specialized equipment, while reliably capturing the complexity of the immune response.

## Introduction

Cellular immune responses are essential for controlling infections by intracellular pathogens, with T cells playing a major role in the induction of an adequate immune response (1). Functional characterization of T cells following infection and vaccination is essential for vaccine development, containment of infection and development of new therapeutic strategies, as highlighted again recently in the ongoing COVID-19 pandemic (1, 2). The important contribution of the T cell response to control human infection is well documented for a broad array of bacterial, fungal and viral etiologies (3–6).

Antigen-specific T cell responses can be evaluated by various methods including Enzyme-linked immunospot (ELISpot) assay (7), intracellular cytokine staining (ICS) in peripheral blood mononuclear cells (PBMC) (8) and whole blood (9, 10), proliferation assay (11), antigen-induced marker (AIM) assay (12) and chromium 51 (^51^Cr) release assay (13). These assays rely on different functional readouts, including cytokine secretion, proliferation, expression of activation markers, and cytotoxicity. Intracellular cytokine staining followed by flow cytometry is one of the most widely used methods for identifying polyfunctionality in T cell subsets by simultaneous detection of multiple cytokines. Studies on cytokine-secreting cells are commonly performed on fresh or cryopreserved PBMC, however both isolation and cryopreservation can negatively impact on cell quality and quantity. In this context, impaired functionality of antigen presenting cells (APC) including monocytes as well as increased apoptosis have been reported (14, 15). Furthermore, some antigen-specific cytokine responses in T cells are reduced following cryopreservation (16). Thus, a whole blood intracellular cytokine assay (WBA) was developed as an alternative (9, 17) and has since been extensively used for the assessment of T cell immune responses in various infectious diseases such as tuberculosis (17, 18), HIV (19), malaria (20) as well as vaccine studies (21).

In this study, we measured T cell responses in patients with scrub typhus, one of the leading causes of undifferentiated febrile illness in endemic tropical regions of Southeast Asia (22). The causative agent *Orientia tsutsugamushi* (OT) is an obligate intracellular Gram-negative bacterium, transmitted by the bite of an infected chigger (larval stage of a mite). A number of studies have shown that the severity of scrub typhus depends on host immune responses (23–26). Results from both animal models and human studies suggest that strong T helper 1 (Th1) responses are associated with acute and severe scrub typhus (27–30). Increased IFN-γ responses are strongly associated with lower levels of bacteremia in a non-human primate model of scrub typhus (31, 32) and a longitudinal study in humans revealed elevated IFN-γ secretion by CD4+ and CD8+ T cells during acute infection (30). In addition, elevated levels of tumor necrosis factor (TNF) correlate with disease severity (33, 34). Of note, all of the aforementioned studies on cytokine-mediated immune responses were assessed in serum, plasma or cryopreserved PBMC.

Characterizing antigen-specific T cell responses in remote areas with limited resources is challenging and requires protocols adapted to the sometimes limited availability of specialized equipment while still capturing the complexity of the immune response. Here we present a WBA protocol specifically optimised for field study sites with respect to ease of setup, duration of antigen stimulation and protein retention. This protocol was designed to assess antigen-specific T cell responses by measuring IFN-γ, TNF and IL-2 production in both CD4+ and CD8+ T cells from whole blood. Using our optimised WBA, we demonstrate polyfunctional CD4+ memory T cell responses to heat-inactivated whole cell OT antigens in humans with acute scrub typhus. In summary, this WBA can be used to answer complex immunological questions in resource-limited settings using low sample volumes, and provides a robust tool to assess T cell responses in infectious disease and vaccine studies.

## Methods

### Ethics Statement

The study was conducted according to the principles of the Declaration of Helsinki (2008) and the International Conference on Harmonization (ICH) Good Clinical Practice (GCP) guidelines. Written informed consent was obtained from all healthy donors and patients enrolled in the study.

The human study protocols were approved by ethics committees of the Faculty of Tropical Medicine, Mahidol University and the Oxford Tropical Research Ethics Committee (FTM-EC), reference number MUTM 2020-021-01. Healthy volunteers were enrolled at Mahidol University under OXTREC reference number 45-14 and used for the optimisation under FTM-EC reference number MUTM 2020-021-01. Patients with scrub typhus were enrolled at Chiangrai Prachanukroh hospital in two scrub typhus studies under OXTREC reference number 46-15 and 4-17.

### Human samples

Venous whole blood from five healthy adult volunteers (age 35-44 yrs) and five scrub typhus patients (age 19-51 yrs) was collected in lithium heparinized tubes. The following inclusion criteria were used for the enrolment of scrub typhus patients: hospitalization with acute undifferentiated fever (temperature > 37.5°C) for ≤14 days and a clinical suspicion of scrub typhus; positive result for scrub typhus using a rapid diagnostic test (ST IgM RDT, InBios International, Seattle, WA, USA or SD BIOLINE Tsutsugamushi Test (IgG, IgM, IgA), Standard Diagnostics ST IgM RDT, Bioline, Korea) or a positive PCR test for OT at Prachanukroh Hospital, Chiangrai.

### *O. tsutsugamushi* antigen preparation

Whole cell antigen *O. tsutsugamushi* (WCA-OT) was prepared as described previously with some adaptations (35). Briefly, the OT strain Karp from Papua New Guinea (Naval Medicine Research Center, NMRC, Silver Spring, Maryland, USA) (36) was propagated in L929 cells (mouse fibroblast cell line, ATCC, CCL-1) cultured in RPMI1640 with L-Glutamine (Gibco, 22400089) supplemented with 10% fetal bovine serum (FBS, Sigma, F7524), in a Biosafety Level 3 (BSL3) containment laboratory. Firstly, L929 cells were grown to 80% confluence and then infected with OT at a multiplicity of infection (MOI) of 10:1 (OT: L929 cell) at 37°C, and 5% CO2 in a humid incubator. At 6 days post infection, the infected cells were harvested by scraping. To release intracellular bacteria L929 cells were disrupted in a bullet blender (Bullet Blender Blue, Nextadvance, USA) at speed setting 8 for 1 min. After centrifugation at 300 g for 3 min to remove cell debris, the supernatant containing the intracellularly grown OT was collected, filtered through a 2.0 μm syringe filter unit and used at MOI 10:1 to infect a fresh preparation of L929 cells (grown to 80% confluence as above). After 5-6 days of culture intracellular bacteria were harvested as described above. The bacteria pellet was washed twice with 0.3 M sucrose solution by centrifugation at 14,000 g for 5 min, subsequently resuspended with sucrose-phosphate-glutamate (SPG) buffer (0.218 M sucrose, 3.76 mM KH2PO4, 7.1 mM KH2PO4, 4.9 mM monosodium L-glutamic acid) and stored at −80°C until ready for heat inactivation. The amount of bacteria was determined by 47 kDa *htra* gene real time PCR assay as described elsewhere (37) using qPCRBIO Probe Master Mix Lo-ROX (PCR Biosystems, PB20.21-05). To find an optimal inactivation method for the antigen preparation which elicits antigen specific T cell responses, WCA-OT was either heat inactivated (HI) or fixed and then tested in a T cell ELISpot assay. To prepare HI-WCA-OT-10^9^ copies/ml of bacterial suspension was made up in R10 media: RPMI 1640 (Sigma, R0883) containing 10% heat-inactivated FBS (Gibco, 10082-147), 2mM L-glutamine (Sigma, G7513) and Penicillin (100 units)-Streptomycin (0.1 mg/ml) (Sigma, P0781) and inactivated for 1 h at 80°C in a water bath. HI-WCA-OT was stored in aliquots at −80°C until further use.

### Whole blood intracellular cytokine assay (WBA)

Whole blood samples were processed within 3 hrs of venipuncture to ensure optimal T cell function and minimize granulocyte activation (38). Stimulation of whole blood (400 μl) was performed in 5 ml round-bottom polystyrene tubes (Falcon, 352054) by adding 100 μl of either HI-WCA-OT at a concentration of 10^9^ OT copies per ml, or staphylococcal enterotoxin B (SEB, Sigma, S4881) at a final concentration of 10 μg/ml as positive control. The co-7 stimulants anti-CD28 (BD Biosciences, 340975) and anti-CD49d (BD Biosciences, 340976) were added at a final concentration of 1 μg/ml each. R10 served as a negative control. WBA optimisation was carried out in the central laboratory (Mahidol-Oxford Tropical Medicine Research Unit, Bangkok, Thailand) using whole blood from healthy human volunteers (n=5). Stimulation was performed using positive and negative controls as described above for 22 or 48 hrs with brefeldin A (BFA, eBioscience, 3615-1) added at a final concentration of 10 μg/ml for the last 4 or 24 hrs of stimulation (see Table 1 for stimulation protocols). In the case of acute scrub typhus samples, whole blood from patients (n=5) was collected at Chiangrai Prachanukroh hospital and processed in a resource-limited laboratory. Samples were stimulated for 22 hrs with BFA added for the last 4 hrs.

**Table 1.** Different stimulation protocols used to optimize the WBA.

| Protocol | Time with antigen before BFA addition (h) | BFA blocking time (h) | Total stimulation time (h) |
| --- | --- | --- | --- |
| SET1 | 18 | 4 | 22 |
| SET2 | 24 | 24 | 48 |
| SET3 | 44 | 4 | 48 |

After the incubation, samples were stained with Near IR Live-Dead fixable dye (Invitrogen, L10119) according to the manufacturer’s instructions. Red blood cell lysis (RBC) was performed by adding 3 ml of 1x FACS lysing solution (BD Biosciences, 349202), followed by vigorous vortexing for 30 sec and incubation for 10 min at room temperature. Cells were then washed twice with 1x FACS lysing solution by centrifugation at 500 g for 5 min, resuspended in 1 ml of freezing media (FBS containing 10% DMSO, Sigma D2650), split into 0.5 ml per tube and frozen in a freezing container at −80°C. In the case of healthy volunteer samples cells were transferred to LN_2_ after 24 hrs. In the case of field study scrub typhus samples, cells were stored at −80°C for a minimum of 2 weeks, then shipped on dry ice to the central laboratory and stored in LN_2_ until further analysis. “Standard Operating Procedure provided in Supplementary Information”

### Flow cytometry staining

Cryopreserved stimulated whole blood samples were slowly thawed in a 37°C water bath, followed by fixation and permeabilization using Cytofix (BD Biosciences, 554722) and Cytoperm/Wash (BD Biosciences, 554723) according to the manufacturer’s instructions. Cells were stained with the following fluorochrome conjugated antibodies to assess polyfunctionality and memory: CD8 V450 (clone RPA-T8, Cat 560347), CD3 V500 (clone SP34-2, Cat 560770), CD4 FITC (clone RPA-T4, Cat 555346 and clone M-T477, Cat 556615), CD45RA-PE (clone HI100, Cat 555489) and IFN-γ APC (clone B27, Cat 554702) all from BD Biosciences as well as IL-2 PE (clone N7.48A, Beckman, Cat IM2718U) and TNF PCP-Cy5.5 (clone MAb11, eBioscience, Cat 45-7349-42). Acquisition of a minimum of 100,000 cells per sample was performed on a MACSQuant 10 analyzer (Miltenyi Biotec). Single-stained controls using mouse Ig/ κ compensation beads (BD Biosciences, 552843) were used to calculate compensation. Flow cytometry data was analyzed using FlowJo™ software v10.5.3 software (BD Life Sciences)(39). Boolean combination gating was performed to assess polyfunctionality of T cells. The gating strategy for flow cytometry analysis of T cell-specific (CD4+ or CD8+) cytokine expression is presented in Supplementary Fig 1.

### Ex-vivo IFN-γ ELISpot assay

To determine IFN-γ secretion by ELISpot, PBMC were isolated from patients with scrub typhus as previously described (29). Briefly, fresh PBMC were added in duplicate wells at 2×10^5^ cells (50 μl) per well and stimulated with 50 μl of HI-WCA-OT (prepared from 10^9^ copies/ml of live OT organism), SEB (final concentration of 10 μg/ml, positive control) and R10 (negative control). A T cell antigen control pool (CEF, Mabtech AB, 3616-1) at a final concentration of 1 μg/ml was used as a source of control antigens. Following 18 hrs of stimulation, IFN-γ secreting cells were detected by Human IFN-γ ELISpot Basic Kit (ALP) (Mabtech AB, 3420-2A). The plates were developed using the AP Conjugate Substrate Kit (Biorad, 1706432) according to the manufacturer’s instructions. ELISpot plates were scanned using a CTL ELISpot reader (Cellular Technology Limited, USA). Spots were counted by Immunospot 3.1 software using the manufacturer’s automated SmartCount™ settings. Results were expressed as IFN-γ spot-forming cells (SFC) per million PBMC.

### Statistical analysis

All results are expressed as median with interquartile range unless otherwise noted. Difference in % live cells and % cytokine secreting cells in the parent population across the three stimulation conditions were analysed by one-way ANOVA followed by Tukey’s multiple comparisons test. Difference in % cytokine secreting cells between HI-WCA-OT stimulated and unstimulated samples were analysed by a nonparametric Mann-Whitney’s (unpaired) t-test. A 2-tailed *p-value* < 0.05 was considered statistically significant. The relationship between the cytokine responses determined by WBA versus ELISpot were tested by Spearman correlation analysis. All statistical analyses and graphical representations were performed using GraphPad Prism version 9.3.0 for Windows (GraphPad Software, San Diego, CA, USA).

## Results

### An overnight stimulation protocol with short Brefeldin A incubation is optimal to assess polyfunctional T cell responses in whole blood

In order to establish an optimal *ex-vivo* stimulation protocol to study antigen-specific T cell responses suitable for field study sites, three different protocols (SET1, SET2 and SET3, Table 1) were compared using heparinized whole blood from healthy donors (n=5) stimulated with staphylococcal enterotoxin B (SEB). SEB was chosen as a positive control due to its ability to induce polyclonal T cell activation with the requirement for APCs and induction of costimulatory signaling via CD28 (40), something that is also necessary for the induction of antigen specific T cell responses.

Two parameters were optimised, the total stimulation time (22 or 48 hrs), and the protein retention time (4 or 24 hrs) using Brefeldin A (BFA) to inhibit the secretion of cytokines. These times were chosen based on data from our previously published protocol for assessment of antigen-specific T cells in PBMC samples (41), and compatibility with the constraints of field study operations including availability of staff and time required to complete patient enrollment, blood sample collection and processing. The expression of the key T cell effector cytokines IFN-γ, TNF and IL-2 was assessed in CD4+ and CD8+ T cell subsets by ICS followed by flow cytometry.

Cell viability was greater than 90% for all stimulation conditions (Fig 1A), with no significant differences between unstimulated and SEB stimulated samples. There was a trend for lower frequency of live cells in SET2, the protocol with the longest BFA incubation time (Fig 1A). To examine the effect of viability staining on data quality, samples were either stained with a fixable live/dead cell stain or not. As expected, in the absence of live/dead cell discrimination the resolution of cell populations was diminished thus confirming the importance of this step in the protocol (see representative density plots in Fig 1B).

**Fig 1.**
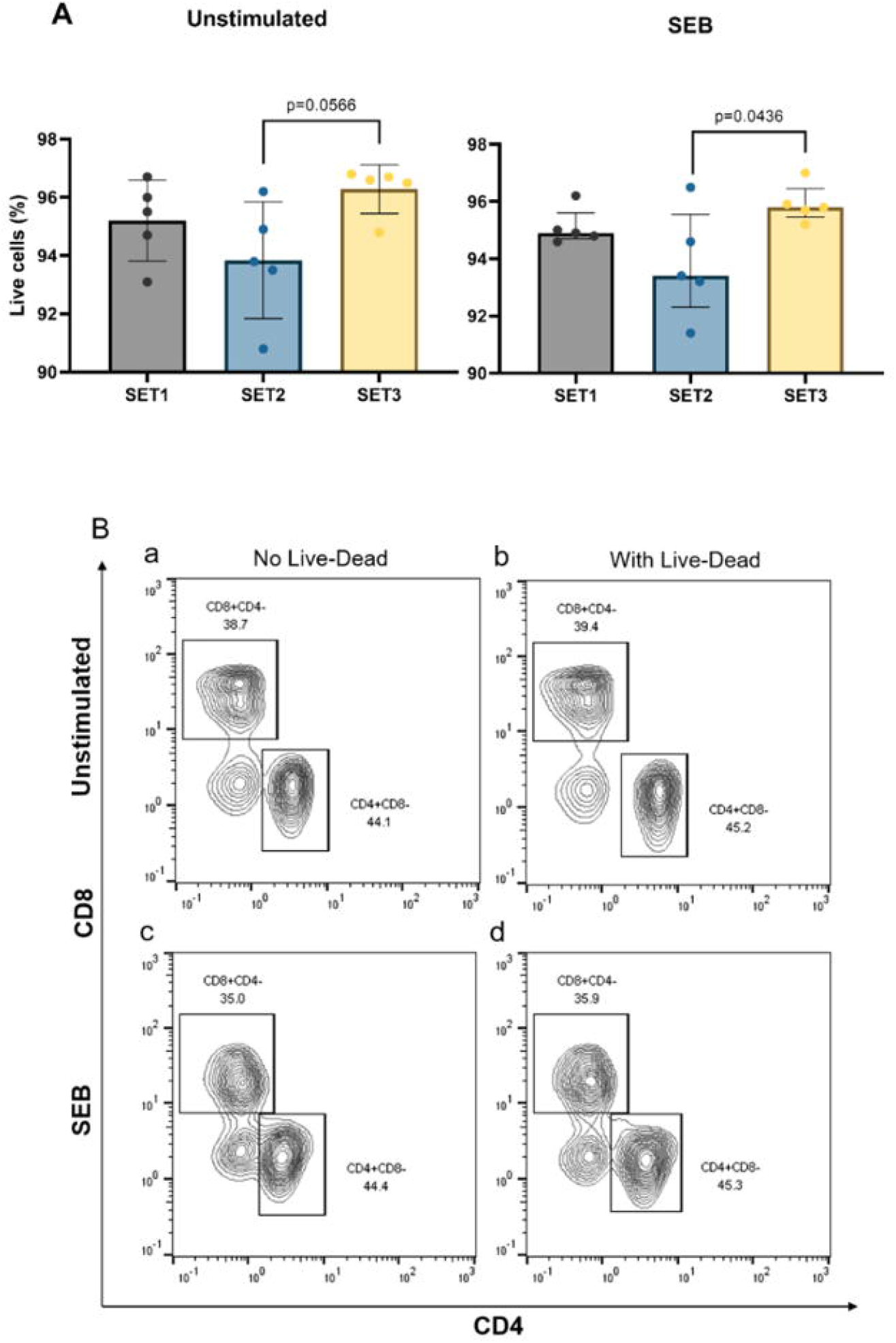
(A) Viability of lymphocytes (% live cells in the single cell gate) in healthy blood samples (Median and IQR for n=5 of biological replicates) stimulated with and without SEB using three different WBA stimulation protocols (SET1, SET2 and SET3, see details in Table 1) followed by ICS and flow cytometry. Difference in % live cells in healthy samples with or without SEB stimulation across the three WBA stimulation protocols were analysed with one-way ANOVA (Kruskal Wallis) followed by Tukey’s multiple comparisons test. Only significant 2-tailed *p* values (p<0.05) are shown in the bar graphs. (B) Representative flow cytometry density plots showing CD4 and CD8 expression on CD3+ T cells stimulated with (c, d) and without SEB (a, b) using protocol SET1 and subsequently stained with (b, d) and without (a, c) a fixable live-dead dye.

The polyfunctionality profile of T cells is considered an important metric reflecting the quality of T cell responses to infection (2). Simultaneous measurement of three T cell effector cytokines allows categorization of cells as single, double or triple cytokine-secreting, and further subdivision based on the specific cytokine combination, resulting in the distinction of 7 different subsets as shown in S1 Table. SET1 and SET2 induced the highest level of total cytokine secretion, with no significant difference between the two protocols for both CD4+ and CD8+ T cells (Fig 2A and B). SET3 with the longest stimulation prior to BFA addition resulted in the lowest overall cytokine levels (Fig 2A and B). Although cytokine responses in SET1 and SET2 were not significantly different, SET2 showed a trend for reduced IFN-γ and IL-2 expression in CD4+ T cells as well as lower frequency of all three cytokines in CD8+ T cells (Fig 2C). Thus, SET1 was chosen as the optimal stimulation protocol.

**Fig 2.**
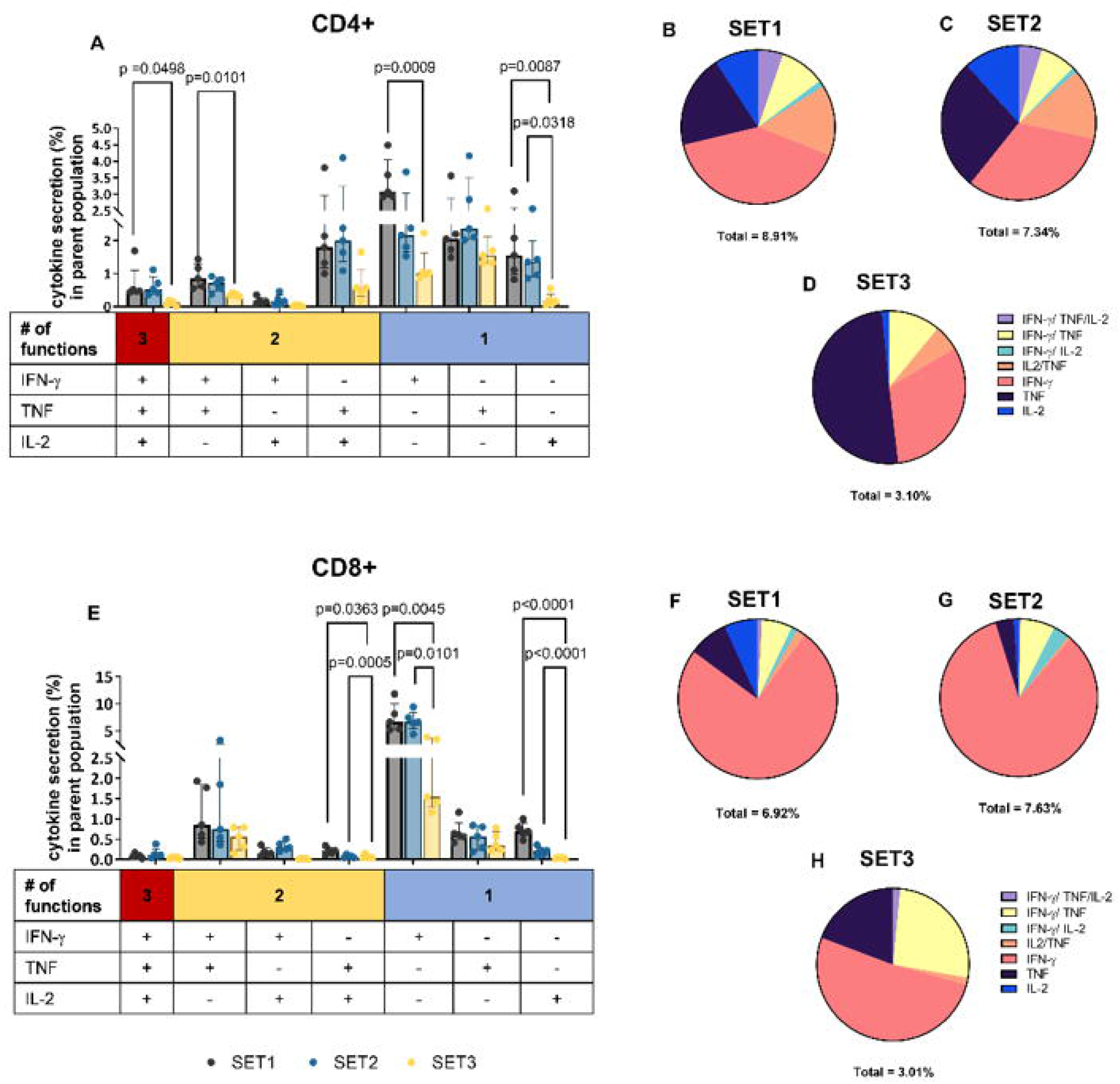
Functional characterization of distinct T cell subsets from healthy donors (Median and IQR for n=5 of biological replicates) using three different WBA stimulation protocols. Frequency of IFN-γ, TNF and IL-2 single (1+), double (2+) or triple (3+) positive CD4+ (A) and CD8+ (E) T cells after stimulation with SEB according to three different protocols (SET1-3) as described in table 1. Difference in % cytokine secreting in parent population in healthy samples stimulated with SEB across the three WBA stimulation protocols were analysed with one-way ANOVA followed by Tukey’s comparisons test, only significant 2-tailed *p* values (p<0.05) are shown on the bar graphs. Pie charts show the proportion of cytokine-producing CD4+ (B-D) and CD8+ (F-H) T cells according to their polyfunctionality. Percentage under each pie chart indicates the proportion of total cytokine secreting cells.

### Antigen-specific polyfunctional Th1 responses are predominant in scrub typhus patients

We next applied the optimised WBA protocol (Fig 3) to study polyfunctional CD4+ and CD8+ T cell immune responses to OT in patients with acute scrub typhus using a heat inactivated whole cell antigen OT preparation (HI-WCA-OT).

**Fig 3.**
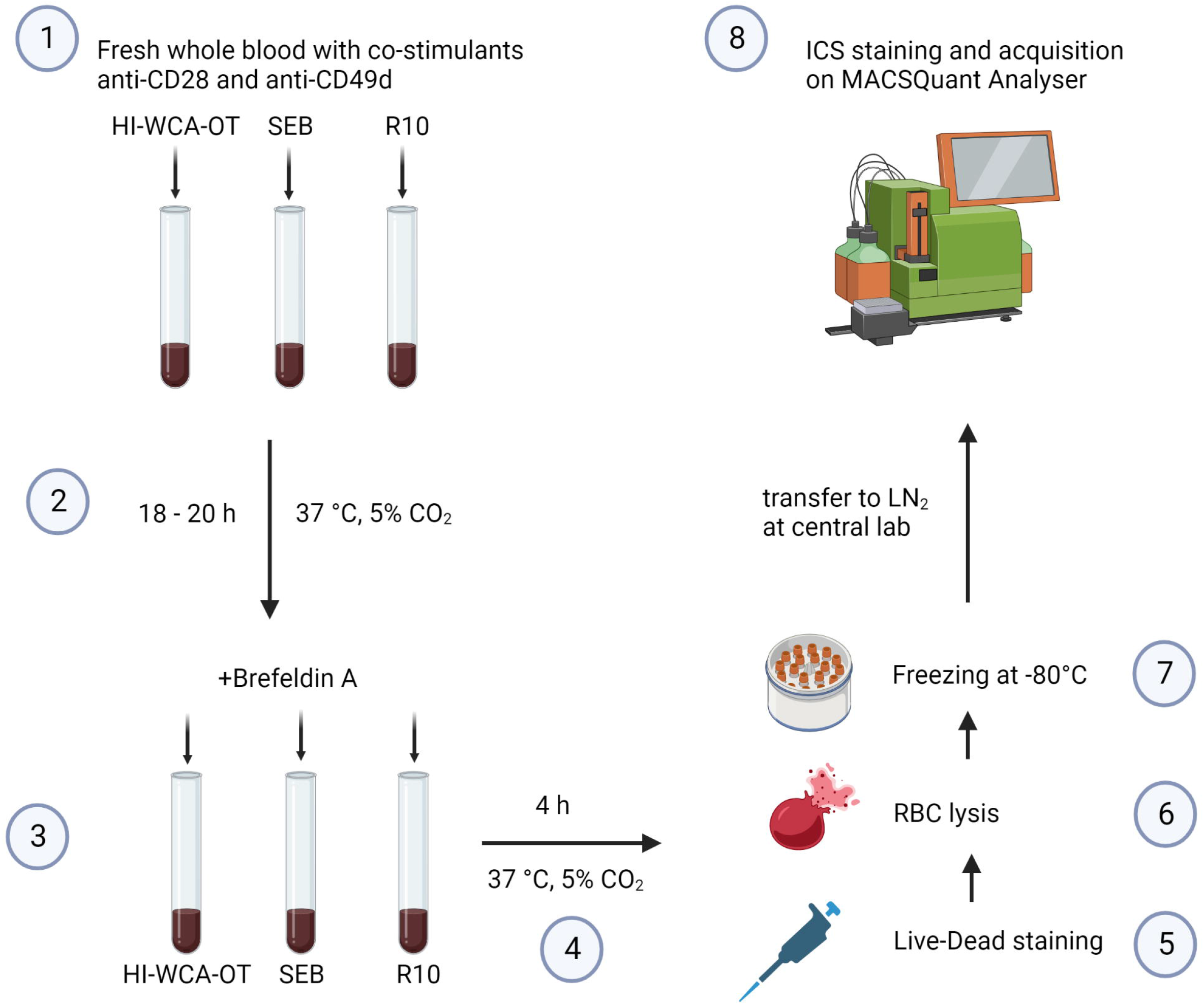
Procedure to perform the optimised WBA for patients with scrub typhus at a field site laboratory. HI-WCA-OT= heat inactivated whole cell antigen *O. tsutsugamushi*, SEB= staphylococcal enterotoxin B, RBC = red blood cell, ICS = intracellular cytokine staining

Analysis of T cell cytokine secretion in response to HI-WCA-OT revealed a predominant CD4+ T cell driven cytokine response, while CD8+ T cell responses were minimal (6.85% of CD4+ T cells versus 0.2% of CD8+ T cells, Fig 4A and B). Following HI-WCA-OT stimulation the majority of cytokine-secreting CD4+ T cells secreted either IFN-γ alone (median=4.15%, range: 2.04 - 13.8%) or were double positive for IFN-γ and TNF (median=1.46%, range: 0.35 - 4.24%) which was significantly upregulated compared to the unstimulated sample (Mann Whitney U-test, p=0.0079). This was followed by secretion of TNF alone (median=0.95%, range: 0.48 - 1.79%). A small proportion of CD4+ T cells were triple positive for IFN-γ, TNF and IL-2 (median=0.15%, range: 0.062 - 0.62 %) which was found to be significantly upregulated compared to the unstimulated sample (p=0.0317) as shown in fig 4A. Interestingly, a high frequency of IFN-γ secreting CD4+ T cells were detected in the unstimulated samples suggesting that the WBA can capture the presence of activated circulating effector T cells in acute scrub typhus. Overall, CD8+ T cells only showed a weak response to OT antigen with a much lower level of cytokine secretion compared to CD4+ T cells. However, similar to CD4+ T cells, the dominant cytokine secreted by CD8+ T cells was IFN-γ (median=0.6%, range: 0.065 - 1.34%), followed by TNF (median=0.18%, range: 0.065 - 0.51%) and both IFN-γ and TNF (median=0.065%, range: 0 - 0.29%) as shown in fig 4C and D. We further used CD45RA expression on CD4+ and CD8+ T cells to determine if the antigen-specific cytokine responses were derived from memory T cells (Fig 5A). Indeed, HI WCA-OT induced IFN-γ secretion almost exclusively from CD45RA negative memory CD4+ T cells and this was significantly upregulated compared to the unstimulated control (Mann Whitney U-test, p=0.0286, Fig 5B).

**Fig 4.**
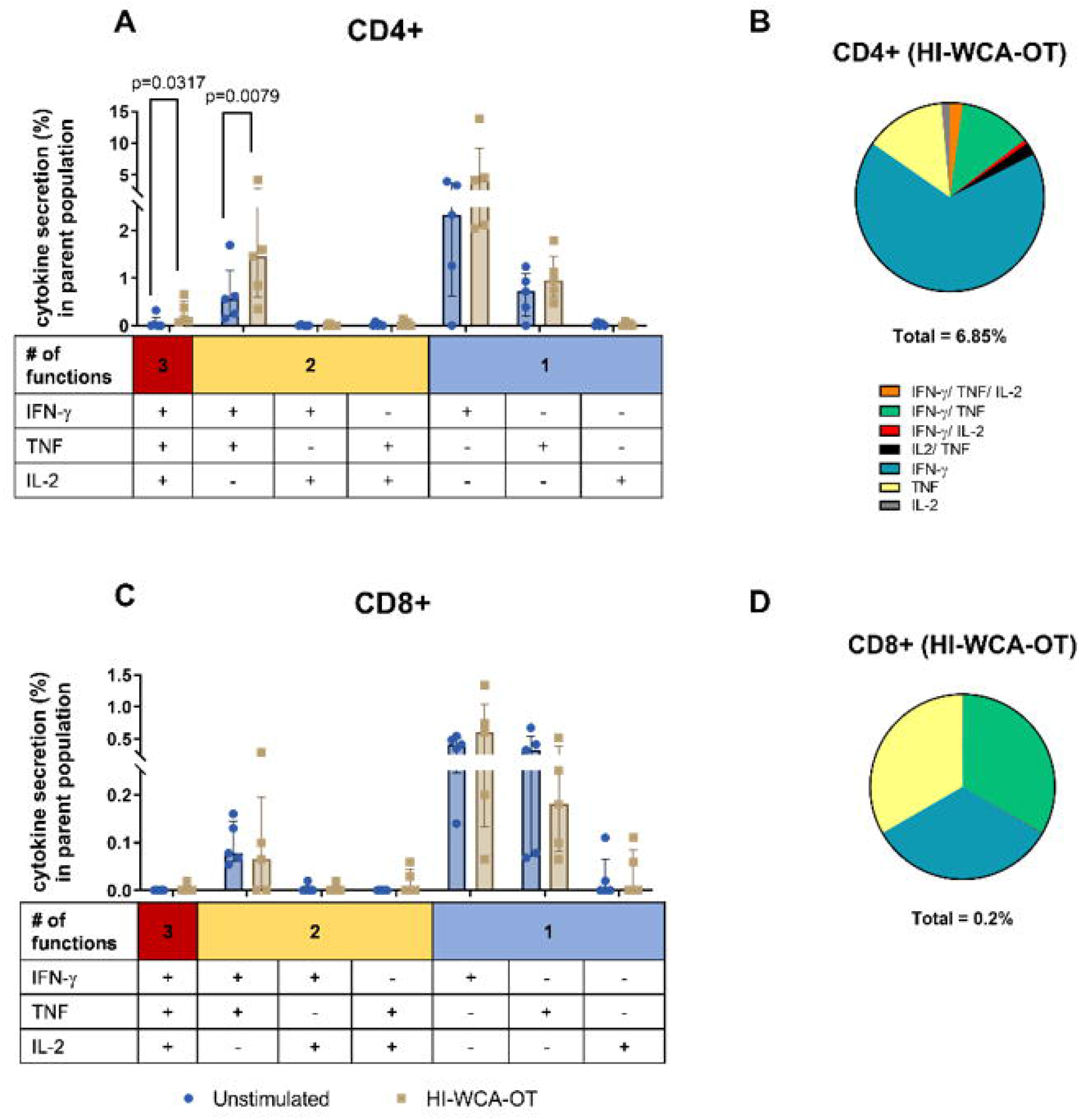
Functional characterization of CD4+ (A) and CD8+ (C) T cell subsets in patients with acute scrub typhus (Median and IQR for n=5 of biological replicates). Frequency of IFN-γ, TNF and IL-2 single (1+), double (2+) or triple (3+) positive CD4+ and CD8+ T cells after stimulation with HI-WCA-OT or without stimulation. Respective pie charts show the proportion of cytokine-producing cells in response to HI-OT-WCA in CD4+ (B) and CD8+ (D) T cells according to their polyfunctionality. Percentage under each pie chart indicates the total frequency of cytokine expressing T cells. Difference in % cytokine secreting in parent population between stimulated and unstimulated samples were analysed by Mann-Whitney U-test, only significant 2-tailed *p* values (p<0.05) are shown in the bar graphs.

**Fig 5.**
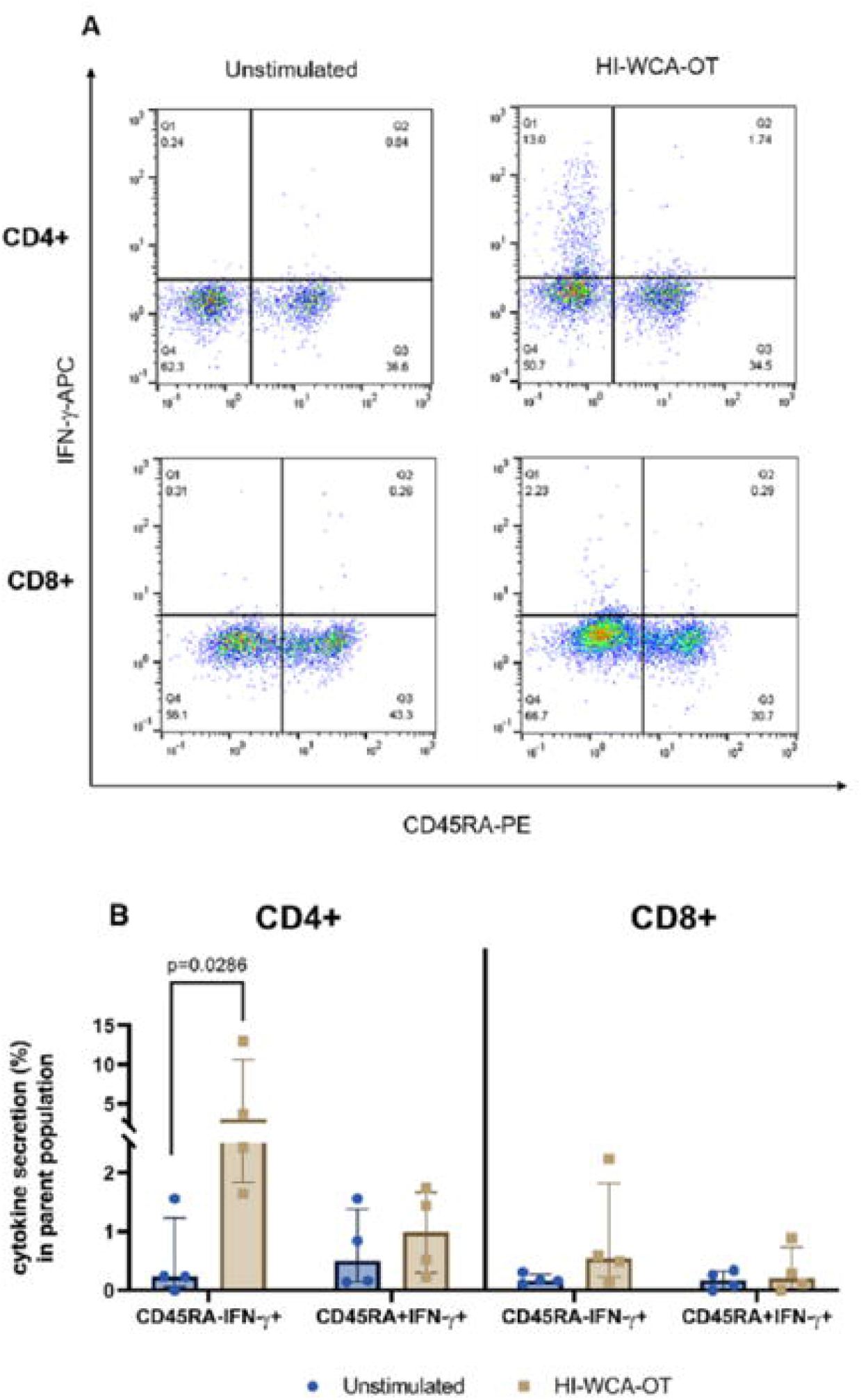
(A) Representative flow cytometry dot plots showing IFN-γ and CD45RA expression on CD4+ and CD8+ T cells following stimulation with HI-WCA-OT or left unstimulated. (B) Frequency of IFN-γ positive CD4 and CD8 T cells amongst CD45RA positive and negative cell subsets when stimulated with HI-WCA-OT and in unstimulated controls (Median IQR for n=4 of biological replicates). Differences in % IFN-γ secreting cells in each subsets between unstimulated and HI-WCA-OT samples were analysed by Mann Whitney U-test. Only significant 2-tailed *p* values (p<0.05) are shown in the bar graph.

In parallel to the WBA, we also performed *ex-vivo* IFN-γ ELISpot assays with freshly isolated PBMC from patients with scrub typhus. The magnitude of the IFN-γ response as measured by IFN-γ ELISpot did not show any significant correlation with IFN-γ secreting CD4+ and CD8+ T cells as assessed by WBA (Spearman r =0.8208, p= 0.1333 for CD4+ and Spearman r =0.4104, p= 0.50 for CD8+) (S2 Fig).

## Discussion

Whole blood is increasingly used as the biological sample of choice to assess antigen-specific T cell function (18, 42, 43). Immediate stimulation of freshly drawn whole blood with antigens of interest avoids compromising cell composition and viability due to excessive handling, and cryopreservation of cells. In addition, it closely mimics the *in vivo* environment by preserving the composition of cells, proteins, antibodies, lipids, metabolites and other blood components. A number of studies have previously shown that the use of whole blood offers significant advantages over the traditional use of PBMC (9, 15, 44–46). Indeed, PBMC isolation and cryopreservation results in a reduction of cellular function, including a decrease in the expression of certain cytokines and surface markers alongside increased apoptosis and cell death (9). Furthermore, monocytes are substantially reduced, thus skewing the monocyte/lymphocyte ratio (9). Storage in liquid nitrogen can also compromise viability and function of certain cell populations including a loss of monocytes (47) and γδ-T cells (48) which are essential for initiating specific cellular immune responses from T cells.

Here we present an optimised WBA for the assessment of cytokine mediated antigen-specific T cell responses in scrub typhus suitable for a field-site laboratory setting.

The induction of cytokine secretion after antigen-specific activation of T cells is a dynamic process depending on a number of factors including antigen concentration, co-stimulatory agent and duration of antigen stimulation (49). ELISpot and ICS assays using fresh or cryopreserved PBMC are widely used to assess antigen-specific T cell responses and determine correlates of protection in infectious disease and vaccine studies. In ELISpot assays, secreted cytokines are captured on a membrane while flow cytometry-based ICS assays capture the intracellular accumulation of cytokines through blocking protein transport. Although the ICS assay is less sensitive than ELISpot it provides valuable qualitative information on magnitude, phenotype and antigen specific function of T cells with the ability to further characterize T cell subsets, memory phenotype and activation. One of the major advantages in using a WBA over the ELISpot assay is that the antigen-specific responses can be assessed shortly after venipuncture, and stimulation occurs in the cell’s natural environment without the need for cell separation and excessive handling. Importantly, the WBA can be performed at resource-limited field sites and stimulated, cryopreserved samples can then be transported to an alternative laboratory for long-term archiving and downstream analyses using specialist equipment. The assay benefits from being relatively scalable using pre-prepared tubes for blood collection containing antigen stimulants, and following centrifugation allowing storage in a freezer until key samples of interest are retrospectively identified for further studies.

The choice of protein transport inhibitor for blocking cytokine secretion is dependent on the cytokine of interest. Two compounds commonly used for this purpose, BFA and monensin, have different cytokine blocking properties (50). We chose BFA for the current study, as previous reports demonstrate superior lymphocyte viability following prolonged stimulation and more efficient intracellular trapping of specific proteins compared to monensin (50).

We tested three different protocols on healthy human blood stimulated with SEB. Higher frequencies of cytokine secreting cells as well as polyfunctional T cells producing IFN-γ, TNF and IL-2 were measured using the protocols with shorter stimulation periods (SET1: 18 hrs and SET2: 24 hrs compared with SET3: 44 hrs) prior to blocking cytokine secretion. Furthermore, we found that longer BFA exposure (SET2: 24 hrs compared with SET1 and 3: 4 hrs) resulted in greater cell death, in agreement with reports demonstrating that prolonged BFA treatment can induce endoplasmic reticulum (ER) stress leading to apoptosis (51). Thus, we chose SET1 (18 hrs + 4 hrs with BFA) as the optimal protocol because it enabled (1) the highest cell viability post stimulation, (2) increased polyfunctionality, (3) higher frequency of T cells secreting a single cytokine of interest (IFN-γ and TNF) and (4) best timeframe to fit into the working routine at the field study site.

To date there is no effective vaccine against scrub typhus and the immune response to the disease has not been fully characterized. By applying the optimised WBA protocol to a clinical study of scrub typhus we demonstrate that the main source of IFN-γ and TNF upon HI-WCA-OT stimulation of WB are CD4+ T cells. Importantly, polyfunctionality (IFN-γ+TNF+IL-2+) of CD4+ T cells was significantly up-regulated as compared to unstimulated cells. Of note, even unstimulated CD4+ and CD8+ T cells showed increased expression of single effector cytokines (IFN-γ and TNF) reflecting the presence of activated effector T cells in peripheral blood of acute scrub typhus patients. Using CD45RA as a marker to distinguish naïve from memory T cells we demonstrate the induction of circulating CD4+ memory responses against OT. Due to the limitations of available channels on our flow cytometer we were not able to further characterize those memory cells. More studies are needed to delineate the phenotype of OT specific memory subsets including markers like CD45RO, CCR7 and CD27.

Our results are consistent with several reports showing strong Th1 responses in acute or severe scrub typhus in both animal models and human (27–30). We observed a trend toward a positive correlation of IFN-γ secreting CD4+ T cells in the WBA and IFN-γ spot forming units in the ELISpot assay, which was not the case for CD8+ T cells, the trend did not achieve a statistical significance, likely due to the small sample size. Several studies using mouse models suggest an important role of CD8+ T cell during scrub typhus (25, 52). Although we detected cytokine responses by CD8+ T cells, they did not show polyfunctional properties. It is possible that the induction of polyfunctional CD8 T cell responses underlies different kinetics in whole blood or predominantly localizes to tissue sites at the time of blood sampling.

Our study has some limitations worth noting. Firstly, a relatively small number of scrub typhus cases was included in the study making it hard to conclude how the WBA compares to the more conventional PBMC based ELISpot assay. Secondly, the assay requires antibodies for co-stimulation and flow cytometry staining making the cost per sample relatively high. Thirdly, the optimized WBA protocol for scrub typhus was only evaluated in one setting. Larger studies in other settings are required to firmly establish the utility of this assay for field study use. One study evaluating human immune responses to melioidosis conducted in North East Thailand ((53) has already successfully adapted this protocol for a different disease thus providing evidence on the versatility and suitability of the WBA protocol for the evaluation of immune responses in rural areas.

## Conclusion

In conclusion, our optimised WBA protocol provides an important analytical tool for the quantification of antigen-specific T cell responses in scrub typhus and can easily be adapted for studying T cell immunity in other infectious diseases and vaccine studies.

## Supporting information

**S1 Table.** Functional T cell-subset classification based on IFN-γ, TNF and IL-2 single, dual and triple cytokine expression

**S1 Fig**. Gating strategy for flow cytometric analysis of T cell (CD4+ or CD8+) cytokine expression. Dead cells were excluded by live-dead cell staining, followed by a single cell gate using forward scatter-area (FSC-A) and height (FSC-H). Lymphocytes were then gated using FSC-A and Side Scatter-area (SSC-A). CD3 positive cells were then selected for further identification of T cell subsets: CD4 or CD8 T positive cells.

**S2 Fig**. Spearman’s correlation analysis of IFN-γ secretion responses measured by WBA and ex-vivo IFN-γ ELISpot assay of CD4+ (A) and CD8+ (B) T cell subsets.

Standard Operating Procedure for Scrub typhus whole blood assay for flow cytometry

## Acknowledgments

We are gratefully thanks to all the participants and staff at Chiangrai Clinical Research Unit (CCRU) at Chiangrai Prachanukroh hospital and Department of Microbiology at MORU for their administration and laboratory support.

## Statements and Declarations

### Disclaimer

“Material has been reviewed by the Walter Reed Army Institute of Research. There is no objection to its presentation and/or publication. The opinions or assertions contained herein are the private views of the author, and are not to be construed as official, or as reflecting true views of the Department of the Army or the Department of Defense. The investigators have adhered to the policies for protection of human subjects as prescribed in AR 70–25.”

### Data availability

“All data reported in the manuscript are publicly available access any data, original software, or materials underpinning the research”

