## Supplementary information for "A whole blood intracellular cytokine assay optimised for field site studies demonstrates polyfunctionality of CD4+ T cells in acute scrub typhus"

**S1 Table.** Functional T cell-subset classification based on IFN-γ, TNF and IL-2 single, dual and triple cytokine expression.

| Functional T cell subset type | Cytokine |
|---|---|
| 1 | IFN-γ |
| 2 | TNF |
| 3 | IL-2 |
| 4 | IFN-γ + TNF |
| 5 | TNF + IL-2 |
| 6 | IFN-γ + IL-2 |
| 7 | IFN-γ + TNF + IL-2 |

**S1 Fig**. Gating strategy for flow cytometric analysis of T cell (CD4+ or CD8+) cytokine expression. Dead cells were excluded by live-dead cell staining, followed by a single cell gate using forward scatter-area (FSC-A) and height (FSC-H). Lymphocytes were then gated using FSC-A and Side Scatter-area (SSC-A). CD3 positive cells were then selected for further identification of T cell subsets: CD4 or CD8 T positive cells.

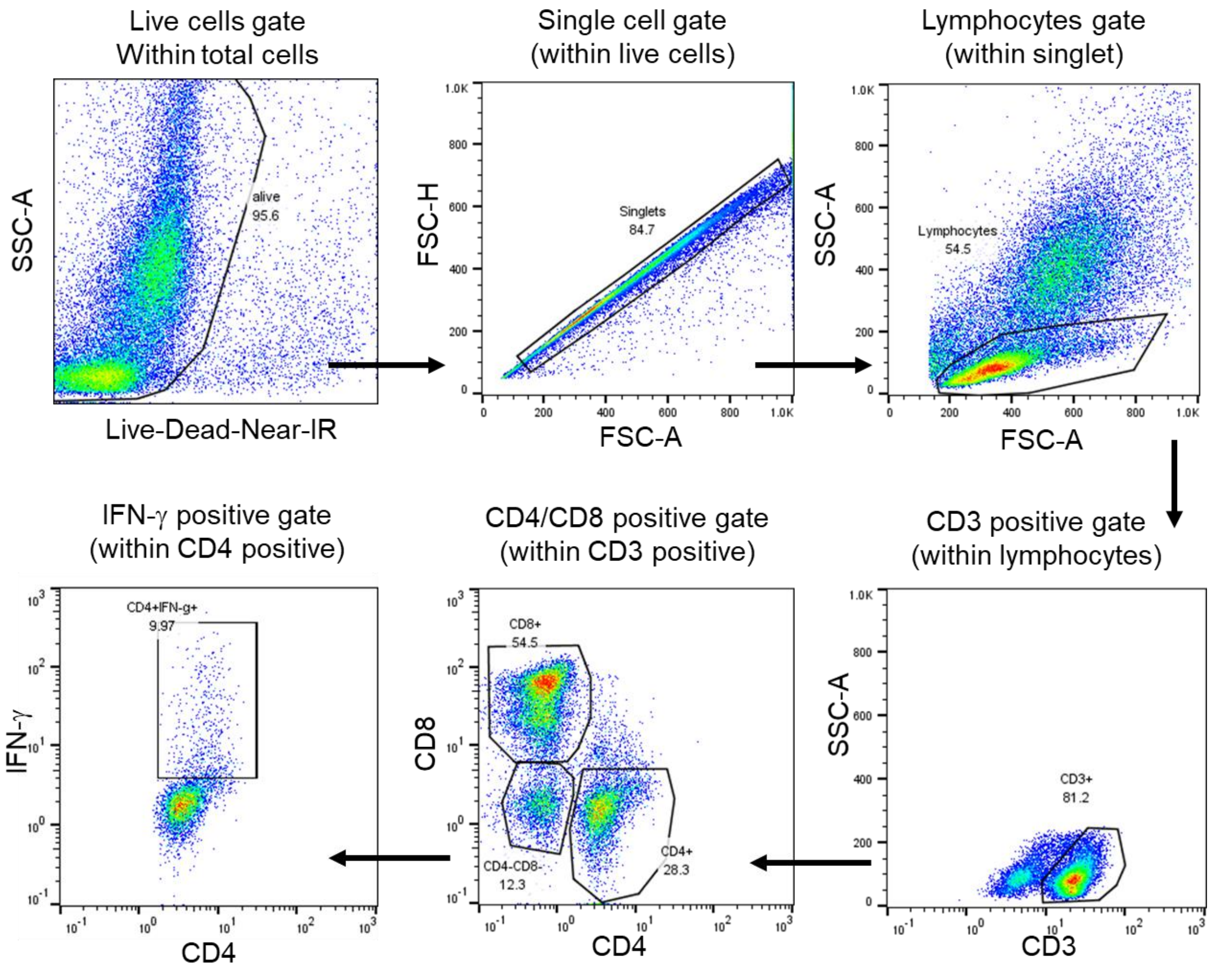

**S2 Fig**. Spearman's correlation analysis of IFN-γ secretion responses measured by WBA and *ex-vivo* IFN-γ ELISpot assay of CD4+ (A) and CD8+ (B) T cell subsets.

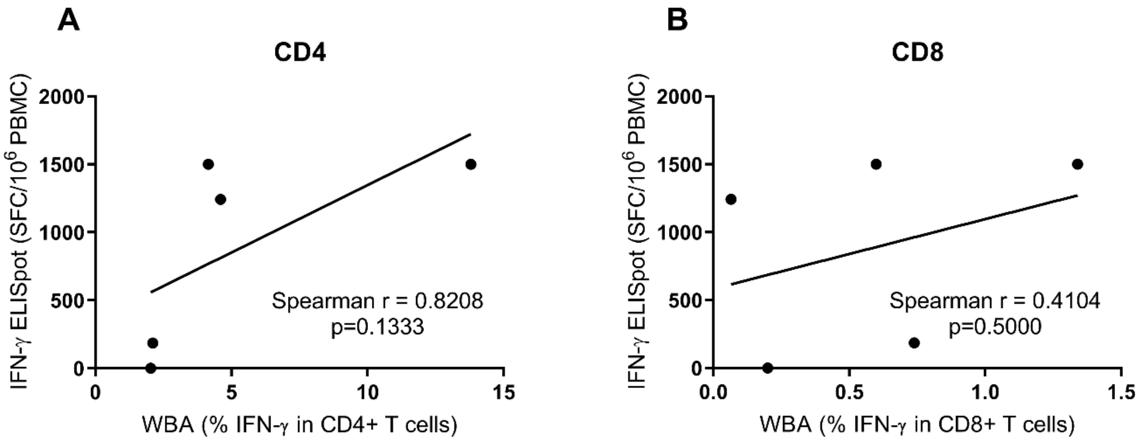

**SOP Title: Whole Blood Assay for Flow Cytometry Scrub typhus**
**Ref: SOP MORU-IM05 version 1.0**
**Owner: Manutsanun Sumonwiriya**
**Review: Biennial**

#### 1.0 Purpose/ Introduction:

Provide a statement of the purpose that this procedure is intended to fulfil.

This protocol provides the procedure for performing a whole blood stimulation assay for intracellular staining

#### 2.0 Scope / Responsibility:

Provide a list as to which specific departments or designees this procedure will apply to.

Laboratory staff working in immunology within the Department of Microbiology, under supervision of Susanna Dunachie

#### 3.0 Definitions:

Provide definitions for all important words and abbreviations referenced in this procedure.

| | |
|---|---|
| DMSO | Dimethylsufoxide, a cryopreservative (protects cells during freezing) |
| FCS | Fetal calf serum |
| ICS | Intracellular staining |
| PBMC | Peripheral blood mononucelar cells |
| PBS | Phosphate buffered saline |
| SEB | Staphylococcal Enterotoxin B |
| PS | Penicillin/streptomycin |
| RPMI | Cell culture medium with bicarbonate buffering system, amino acids and vitamins |
| RT | Room temperature |

#### 4.0 Associated Standard Forms:

List all forms associated with this procedure. none

#### 5.0 Document Change History

This section is to be completed by Quality Management or designee

| Date: | Description of Change: | Initiator: | Reviewer: | Approver: |
|---|---|---|---|---|
| 1st May 2015 | First draft | MS | | |
| 10thJul 2015 | Antigens for stimulation, change from proteins to peptide megapools | MS | | |
| 05th Aug 2015 | Concentration of peptide megapools | MS | | |
| 27th June 2017 | Change antigens panel | MS | | |
| 23rd Aug 2017 | Add Live-Dead Staining | MS | | |

#### 6.0 Responsibilities

The senior immunologist is responsible for supervising the use of this protocol by laboratory staff, in accordance with biosafety requirements. **This protocol uses human/animal blood. All users MUST have health and safety clearance**

#### 7.0 The Procedure

Samples

Collect fresh whole blood in Li-heparinized tubes and use in assay within 3 hours post blood draw. A total of 2.5ml whole blood are needed to perform the WBA described below using 5 different stimulation conditions.

Solutions to be made up

R0

- 490 ml RPMI 1640 (without glutamine)
- 5 ml 10000 U/ml penicillin/streptomycin from aliquot.
- 5 ml 200 mM L-Glutamine from aliquot.
- Label “R0” and write date, initials, and write PS and LG lot numbers on the bottle. Store at 4°C and replace after 1 month or if any concern about sterility

PBS

- Made with PBS tablets from SIGMA, and dH2O. 1 tablet in 100ml water.
- Autoclave

| Reagent: | Catalog No. | Stock solution: | Working Dilution: |
|---|---|---|---|
| **"COCKTAIL"**<br>**Anti-CD28**<br>**Anti-CD49d** | BD Bioscience<br>340975<br>340976 | Each provided as 200 µg in 0.01% azide in PBS; **1** mg/mL. Store at 4°C in the dark. | Dilute 1: 10 of each in a 1 ml tube, combine<br>3 µl anti-CD28<br>3 µl anti-CD49d<br>24 µl PBS. Label "**COCKTAIL**"<br>For 500 µL blood; add 5 µl of the above "**COCKTAIL**" solution for a final concentration of 1 µg/mL each.<br>For Full antigen panel (=5 Ags) a total of 25 µl of the cocktail are needed per sample |
| "**Heat Inactivated WCA-OT-Karp**"<br>Whole killed *Orientia tsutsugamushi* | From In-house propagation | Working WCA-OT is $\mathbf{10^9}$ copies/ml (Aliquots stored at -80°C Freezer. Keep at 4°C once thawed) | Add 100 µl of this $\mathbf{10^9}$ copies/ml "**Live-WCA-OT-Karp**" to tube for final conc of $10^8$ copies/tube in 400ul of blood |
| **r56-UT76 peptide megapool** | Mimotope<br>(New lot#1/2017) | Stock r56 peptide megapool is **20** ug/ml. Aliquots stored at -80°C. Keep at 4°C once thawed | Add 100 µl of this 20 µg/ml "**r56-UT76 peptide megapool**" to tube for final conc of 4 µg/ml in 400 µl of blood |
| **r47-UT76 peptide megapool** | Mimotope<br>(New lot#1/2017) | Stock r47 peptide megapool is **20** µg/ml. Aliquots stored at -80°C. Keep at 4°C once thawed | Add 100 µl of this 20 µg/ml "**r47-UT76 peptide megapool**" to tube for final conc of 4 µg/ml in 400 µl of blood |
| **"SEB WBA"** | Sigma | Stock SEB is 1 mg/ml. Keep at 4°C once thawed | Dilute 1: 20 of stock SEB (1mg/ml) in R10<br>Add 100 µl of this "**SEB WBA**" to each tube for final conc of 10 µg/ml in 400 µl blood |
| **Brefeldin-A** | Ebioscience<br>00-4506-51 | Stock 3 mg/ml in methanol | Dilute stock 1:15 in sterile PBS to 200 µg/ml by adding 20 µL stock BfA + 280 µL PBS); do not store. To **500** µL of blood, add 25 µL for a final concentration of 10 µg/ml. For Full antigen panel (=5 Ags) 125 µl of the cocktail are needed per sample |
| **Cryosolution** | DMSO: Sigma D2650 | 10% DMSO in heat-inactivated FCS, store at 4C in dark | |
| **FACS Lysing Solution** | BD Bioscience<br>349202 | Provided as 100mL of a10X solution. Store at ROOM TEMPERATURE | Dilute 1 part 10X FACS Lysing Solution in 9 parts ROOM TEMPERATURE sterile WATER (not PBS). May store this at RT for future use. To 500uL blood, add 3 mL FACS Lysing Solution. |
| **Near IR Live-Dead fixable dye** | Invitrogen L10119 | Add 50 µl DMSO (provided with the kit) to make a 1mM stock | Add 1 µl of 1 mM dye to the sample tube (~ 500 µl o stimulated blood) |

#### Hardware:

| Product: | Catalog No. | Comments: |
|---|---|---|
| 5.0mL falcon BD tubes | | Use the type with caps |
| $CO_2$ incubator | | 37°C |
| Benchtop Centrifuge | | Use with swing bucket rotor or use a water bath as an alternative option |

### Procedure (continue)

**Please refer to the following reference for more info, and for flow cytometry steps in the assay:**
Hanekom WA, Hughes J, Mavinkurve M, Mendillo M, Gelderbloem SJ, Watkins M, Gamieldien H, Sidibana M, Davids V, Mansoor N, Hawkridge A, Haslett PAJ, Ress S, Hussey GD, Kaplan G. Novel application of a whole blood intracellular cytokine detection assay for evaluating specific T cell frequency and function in field studies. Journal of Immunologic Methods 2004; 291:185-195.

STIMULATION:

1. Each subject in the study needs 5 x 5 ml tubes labelled with their ID number and stimulant (antigen or control) i.e. "HI-WCA-OT" "r56", "r47", "SEB" or "R0"
2. Prepare pre-labelled 5 ml tubes to each contain **5** µl of "**COCKTAIL**" (co-stimulant) and **NO Bfa.** Add the stimulants as follows to respective tubes:
    a. **Heat Inactivated-WCA-OT-Karp: Add 100 µl of working "HI-WCA-OT-Karp"**
    b. **r56: Add 100 µl of working "r56 megapool"**
    c. **r47: Add 100 µl of working "r47 megapool"**
    d. **SEB: Add 100 µl of working "SEB WBA"**
    e. **0: No antigen, do not add anything**
3. Add **400uL** (<u>Li-heparinized NOT other anticoagulants!</u>) to each tube, cap tightly, vortex.
4. Incubate at 37ºC for **18-20** hours in $CO_2$ incubator or in a water bath (optional).
5. At **18-20** hours after incubation, add **25** µL Bfa (10ug/mL). Incubate again for a further **4** hours.

HARVEST:

**Washing in PBS may help improve lysis

6. At harvest time, stain with Live-Dead Staining dye (Near IR) 1 µl which enables discrimination of live from dead cells in subsequent flow cytometry analysis.
7. Incubate on ice for 20 min.
8. Add PBS 1 ml/tube for washing then centrifugation at 500 rcf for 5 min at RT.
9. Remove supernatant by carefully pipetting without disturbing the pellet (RBC)
10. Add **3 ml** of FACS Lysing solution to lyse RBC and vortex.
11. **Incubate at room temperature for 10 min.** (to help RBC lysis), vortex for 30 s.
12. Spin in centrifuge for 5 min at 500 rcf
13. Remove supernatant by pouring off then vortex to resuspend pellet for 10 s
14. Repeat RBC lysis steps 10-12
15. Remove supernatant by pouring off then vortex to resuspend pellet
16. Add 1ml freezing mix to resuspended pellet and transfer 500ul each into 2 labelled cryotubes (so that there is two tubes for each condition for freezing) followed by stepwise freezing to -80°C

#### 8.0 Risk assessment

| COSHH risk assessment - University of Oxford COSHH Assessment Form | |
|---|---|
| **Description of procedure**<br>Cell stimulation | **Substances used**<br>BD Biosciences antibodies for cell stimulation |
| **Quantities used**<br>microlitres | **Frequency of use**<br>Less than monthly |
| **Hazards identified**<br>0.01% sodium azide – potential hazard if swallowed, inhaled or skin contact of significant quantity | **Could a less hazardous substance be used instead?**<br>NO |
| **What measures have you taken to control risk?**<br>Good laboratory practice, including use of gloves and PPE<br>Working within BSC if possible, avoid creating aerosols | |
| **Checks on control measures**<br>Observation and supervision by senior staff | |
| **Is health surveillance required?**<br>No | **Training requirements:**<br>GLP |
| **Emergency procedures**:<br>Report all incidents to Safety Adviser<br>Eye wash for splashes | **Waste disposal procedures**:<br>Autoclaving of materials used |
